# Gene expression responses of CF airway epithelial cells exposed to elexacaftor/tezacaftor/ivacaftor (ETI) suggest benefits beyond improved CFTR channel function

**DOI:** 10.1101/2024.08.28.610162

**Authors:** Thomas H. Hampton, Roxanna Barnaby, Carolyn Roche, Amanda Nymon, Kiyoshi Ferreira Fukutani, Todd A. MacKenzie, Bruce A. Stanton

## Abstract

The combination of elexacaftor/tezacaftor/ivacaftor (ETI, Trikafta) reverses the primary defect in Cystic Fibrosis (CF) by improving CFTR mediated Cl^-^ and HCO_3_^-^ secretion by airway epithelial cells (AEC), leading to improved lung function and less frequent exacerbations and hospitalizations. However, studies have shown that CFTR modulators like ivacaftor, a component of ETI, has numerous effects on CF cells beyond improved CFTR channel function. Because little is known about the effect of ETI on CF AEC gene expression we exposed primary human AEC to ETI for 48 hours and interrogated the transcriptome by RNA-seq and qPCR. ETI increased defensin gene expression (*DEFB1*) an observation consistent with reports of decreased bacterial burden in the lungs of people with CF (pwCF). ETI also decreased *MMP10* and *MMP12* gene expression, suggesting that ETI may reduce proteolytic induced lung destruction in CF. ETI also reduced the expression of the stress response gene heme oxygenase (*HMOX1*). qPCR analysis confirmed *DEFB1, HMOX1, MMP10* and *MMP12* gene expression results observed by RNA-seq. Gene pathway analysis revealed that ETI decreased inflammatory signaling, cellular proliferation and MHC Class II antigen presentation. Collectively, these findings suggest that the clinical observation that ETI reduces lung infections in pwCF is related in part to drug induced increases in *DEFB1*, and that ETI may reduce lung damage by reducing *MMP10* and *MMP12* gene expression, which is predicted to reduce matrix metalloprotease activity. Moreover, pathway analysis also identified several genes responsible for the ETI induced reduction in inflammation observed in people with CF.

**New and Noteworthy:** Gene expression responses by CF AEC exposed to ETI suggest that in addition to improving CFTR channel function, ETI is likely to increase resistance to bacterial infection by increasing levels of beta defensin 1 (hBD-1). ETI may also reduce lung damage by suppressing MMP10, and reduce airway inflammation by repressing proinflammatory cytokine secretion by AEC cells.

## Introduction

Cystic Fibrosis (CF) is a recessive genetic disease caused by mutations in the gene that codes for the cystic fibrosis transmembrane conductance regulator, CFTR (1). Mutations in the *CFTR* gene lead to chronic bacterial lung infections, prolonged and excessive inflammation and a progressive decrease in lung function (1, 2). Several drugs have been developed for people with CF (pwCF), including Kalydeco^®^, Orkambi^®^ and Symdeko^®^ (3). Kalydeco^®^ (ivacaftor) improves CFTR function and clinical outcomes for class III channel gating mutations (e.g., G551D), Orkambi^®^, the combination of ivacaftor and lumacaftor (4), improves CFTR function and clinical outcomes of people homozygous for the ΔF508 mutation, and Symdeko^®^ (tezacaftor and ivacaftor) improves CFTR function and clinical outcomes of people homozygous for the ΔF508 mutation, and for pwCF with one copy of ΔF508 and a mutation in CFTR responsive to Symdeko^®^ (5). Trikafta^®^ (<u>e</u>lexacaftor, tezacaftor and ivacaftor-ETI), approved by the FDA in 2019, is approved for pwCF having at least one copy of the ΔF508 mutation, the most common mutation that causes CF (6). In clinical trials Trikafta^®^ has been shown to be more effective than Orkambi^®^ and Symdeko^®^ in improving lung function, and reducing exacerbations and hospitalizations (6–9). Although the effects of ETI on Cl^-^ secretion by mutant CFTR have been investigated (10), there are no reports describing the effect of ETI on comprehensive gene expression by CF AEC. Thus, the goal of studies in this report are to increase our knowledge of the effects of ETI on gene expression in human primary CF AEC.

Gene expression studies of blood samples drawn from pwCF have demonstrated that ivacaftor induces gene expression patterns associated with a more robust immune response (11), that subjects with the largest gene expression responses to ivacaftor experience the greatest clinical benefit (12), and that the combination of ivacaftor and lumacaftor decreased the expression of cell-death genes, *MMP9* and *SOCS3* (13). Although transcriptional responses in blood are relevant to CF, they do not capture responses in CF lung epithelial cells, and much less is known about how lung epithelial cells respond to modulator therapy at the transcriptional level. A transcriptional study of the IB3-1 CF line exposed to ivacaftor or lumacaftor reported no significant impact on immune gene signaling (14) but single cell transcriptomic analysis of nasal swabs in children with CF showed that ETI partly restored interferon signaling in epithelial cells to levels seen in controls (15). A recent study of CF and wild type airway epithelial exposed to ETI for 72 hours reported many transcriptional differences between CF and wild type cells but no response to ETI (16). Nasal epithelial transcriptomic profiles have recently been shown to predict changes in FEV_1_ (forced expiratory volume in 1 second) and BMI (body mass index) with ETI treatment (17). However, little is known about the response of CF AEC to ETI. An improved understanding of ETI-induced changes in gene expression in CF AEC will provide important information for future drug development efforts in CF.

To address this knowledge gap, we exposed human primary AEC from five CF donors in air liquid interface culture to ETI for 48h and measured gene expression by RNA-seq as described in Methods. We found that 11 genes were differentially expressed by ETI (FDR < .05), including *DEFB1, HMOX1, MMP10*, and *MMP12*. Transcriptional changes in gene expression were validated by qPCR. Gene pathway analysis revealed that ETI decreased inflammatory signaling, cellular proliferation and MHC Class II antigen presentation. Collectively, the data suggest that the ETI induced reduction in lung infections in pwCF are related in part to drug induced increases in *DEFB1*, and that ETI may reduce lung damage by reducing *MMP10* and *MMP12* gene expression, which is predicted to reduce matrix metalloprotease activity. Moreover, pathway analysis also identified several genes responsible for the ETI induced reduction in inflammation observed in people with CF.

## Methods

### Cell culture and ETI exposure

De-identified primary human AECs from five CF donors homozygous for the ΔF508 mutation were obtained from Dr. Scott Randell (University of North Carolina at Chapel Hill, Chapel Hill, NC) and grown in culture as previously described (18, 19). One of the donors was male and four of the donors were female. Donors were between 14 and 27 years old. In brief, cells were grown in BronchiaLife basal medium (Lifeline Cell Technology, Frederick, MD, Cat. #LM-0007) supplemented with the BronchiaLife B/T LifeFactors Kit (Lifeline Cell Technology, Cat. #LS-1047) and penicillin (10,000 U/mL) and streptomycin (10,000 μg/mL) (Sigma-Aldrich, Cat. #P4333). Cells were studied between passages 4–9 and results were independent of passage number. The Dartmouth Committee for the Protection of Human Subjects determined that the use of CF AEC cells in this report is not considered human subject research because cells are taken from discarded tissue and contain no patient identifiers. 500,000 CF AEC were seeded on 24-mm permeable supports (Corning, Corning, NY, Cat. #3407), coated with Collagen type IV (Sigma-Aldrich, Cat #C-7521) and grown in an air-liquid interface media (ALI) (University of North Carolina, Chapel Hill, NC) at 37°C for 3 to 4 weeks to establish polarized monolayers (20, 21). Ivacaftor (10 nM) (Selleck Chemicals, Houston, TX, Cat. #S1144), <u>e</u>lexacaftor (3 μM) (Selleck Chemicals, Houston, TX, Cat. #S8851), and tezacaftor (3 μM) (Selleck Chemicals, Houston, Tx, Cat. #S7059) or vehicle (DMSO) (Fisher Scientific, Hampton, NH, Cat. #D128-500) were added to the basolateral media 48 h before experiments. ETI was added for 24 hours, media was removed and fresh media with ETI was added back for an additional 24 hours. The total exposure to ETI was 48 hours.

After 48 hours of exposure to ETI or vehicle (DMSO), polarized CF AEC were rinsed three times with PBS ++ (Invitrogen, Waltham, MA, Cat. #14190-250), and RNA was isolated by the miRNeasy kit (Qiagen, Germantown, MD, Cat. #217004). cDNA synthesis was performed using RNA (1 μg) as input using SuperScript IV (Invitrogen, Grand Island, NY, Cat. #18090050). qPCR was performed using TaqMan Master Mix (Invitrogen, Waltham, MA, Cat. # 4304437) as recommended by the manufacturer.

### RNA-seq

Total RNA was isolated using the miRNeasy kit (Qiagen, Germantown, MD, Cat. #217004). RNA for RNA-seq was quantified by Qubit and integrity measured on a fragment analyzer (Agilent). 100 ng RNA was used as input into the Quantseq FWD workflow (Lexogen) for library preparation following manufacturer’s instructions. Libraries were pooled for sequencing on a NextSeq2000 instrument targeting 10M, single-end 100 bp reads per sample. Paired-end read were trimmed using cutadapt v4.0 (22), and aligned to the human genome reference GRCh38, using Hisat2 v2.1.0 (23). Genes were quantified using FeatureCounts v2.0.1 (24), with Human Ensembl v97 as the gene annotation reference. Alignment and quantification metrics were calculated using Samtools v1.15.1 (25).

### Filtering and Normalization

Filtering and normalization were performed in edgeR version 4.0.1 (26) as follows. Genes with fewer than 10 counts in any given sample or fewer than 15 counts across all 10 samples were filtered out using the filterByExpr function, leaving 15,229 genes for downstream analysis. Library sizes were adjusted using calcNormFactors and normalized log base 2 expression values were retrieved using the cpm function in edgeR.

### Exploratory data analysis

Euclidean distances between samples were calculated using the dist function in base R and an adonis test was performed using adonis2 in the R package vegan (27) and visualized using fviz_pca_ind in package factoextra 1.0.7. (28).

### Differential gene expression

Differential gene expression analysis of RNA-seq data was performed using genewise negative binomial generalized linear models in edgeR version 4.0.1 (26). Fastq files for all RNA-seq samples as well as count tables of raw and normalized aligned reads have been deposited in NCBI’s Gene Expression Omnibus (29) GSE268718 (https://www.ncbi.nlm.nih.gov/geo/query/acc.cgi?acc=GSE268718).

### Ingenuity Pathway Analysis

Significantly activated or repressed gene pathways were identified using QIAGEN IPA (QIAGEN Inc., https://digitalinsights.qiagen.com/IPA) as follows. ENSEMBL gene identifiers, log base 2 fold change responses to ETI compared to DMSO, and unadjusted P values (https://figshare.com/articles/dataset/ETI_edgeR_Results/25975324) were uploaded to IPA and mapped to the Ingenuity knowledge base, and a new core analysis was created. Significantly differentially expressed genes for this analysis were defined as those with an absolute log2 fold change > 1 and an unadjusted P value < .05, resulting in 281 analysis-ready genes of which 178 were repressed and 103 were induced. Fisher’s exact tests of gene set enrichment performed by IPA on 537 Ingenuity canonical pathways identified 7 pathways with an absolute activation z score > 1 and a P value < .05 and relevant to airway epithelial cells.

### Measurement of gene expression by qPCR

Quantitative polymerase chain reaction (qPCR) was used to confirm significant ETI effects observed in RNA-seq analysis for *HMOX1, DEFB1, MMP10* and *MMP12*, and to explore the possibility that qPCR might detect significant changes in *IL1B* and *TNF*, two genes not significantly differentially expressed (FDR < .05) in RNA-seq, but relevant to the airway immune response in CF (30, 31). In addition, ETI responses were measured by qPCR in candidate reference genes *GAPDH, GUSB, HPRT1, HSP90AB* and *UBC*.

Triplicate measurements of cycle threshold (CT) of a given gene on the same PCR plate were averaged to create a matrix of CT values for each target and reference gene in each condition (https://figshare.com/articles/dataset/Cycle_threshold_values_of_PCR_measurements_for_target_and_reference_genes_/25997890). The three most stable reference genes (*HSP90AB, GAPDH, HPRT1*) were identified by the geNorm2 function in the R package ctrlGene (32). Mixed effect linear models in lme4_1.1-35.3 (33) and P values were generated using lmerTest_3.1-3 (34) to estimate the response to ETI, using the average CT of *HSP90AB, GAPDH, HPRT1* to control for differences in sample quality unrelated to treatment, modeling airway cell donor as a random effect (18).

### Measurement of Cytokines and Beta Defensin 1 by ELISA

Secretion of hBD-1 (PeproTech: #900-K202), IL-6, IL-8, G-CSF, and MCP1 (R & D Systems: #DY206, #DY208, #DY214, #DY279) by CF AEC exposed to ETI or vehicle was measured by ELISA in triplicate wells per sample as recommended by R & D Systems. Concentrations of analytes measured by ELISA were calculated from standard curves of concentration as a function of fluorescence fitted with a four parameter model using the R drm function in the drc 3.0-1 package (35). The statistical significance of ETI effects was assessed based on one tailed t tests of log base2 fold changes. The alternative hypothesis of these tests was contingent on the direction of change of the corresponding gene measured by RNA-seq. For example, if the gene corresponding to beta defensin 1 (*DEFB1*) was induced, then the alternative hypothesis for the test was “greater.”

### General statistical analysis

Data were analyzed using the R statistical platform (36). Figures were created using ggplot2 (37). The figure legends describe the specifics of the statistical analysis of each data set.

## Results

### Differences between donors explain most of the divergence between control and ETI exposure

Human primary AEC cells in ALI culture obtained from five donors with CF were exposed to ETI or vehicle for 48 h. RNA was isolated, cDNA libraries were created, sequenced, aligned to a human reference genome, and normalized in edgeR as described in Methods. Dimension reduction by Principal Components Analysis (Figure 1A) revealed that samples clustered closer together by donor rather than ETI treatment. An adonis test of Euclidean distances between the 10 samples verified that 70% of the variability between samples can be explained by differences between donors (P = 0.001) and only 6% of differences were explained by ETI treatment, although this effect did not reach significance (P = 0.35).

**Figure 1.**
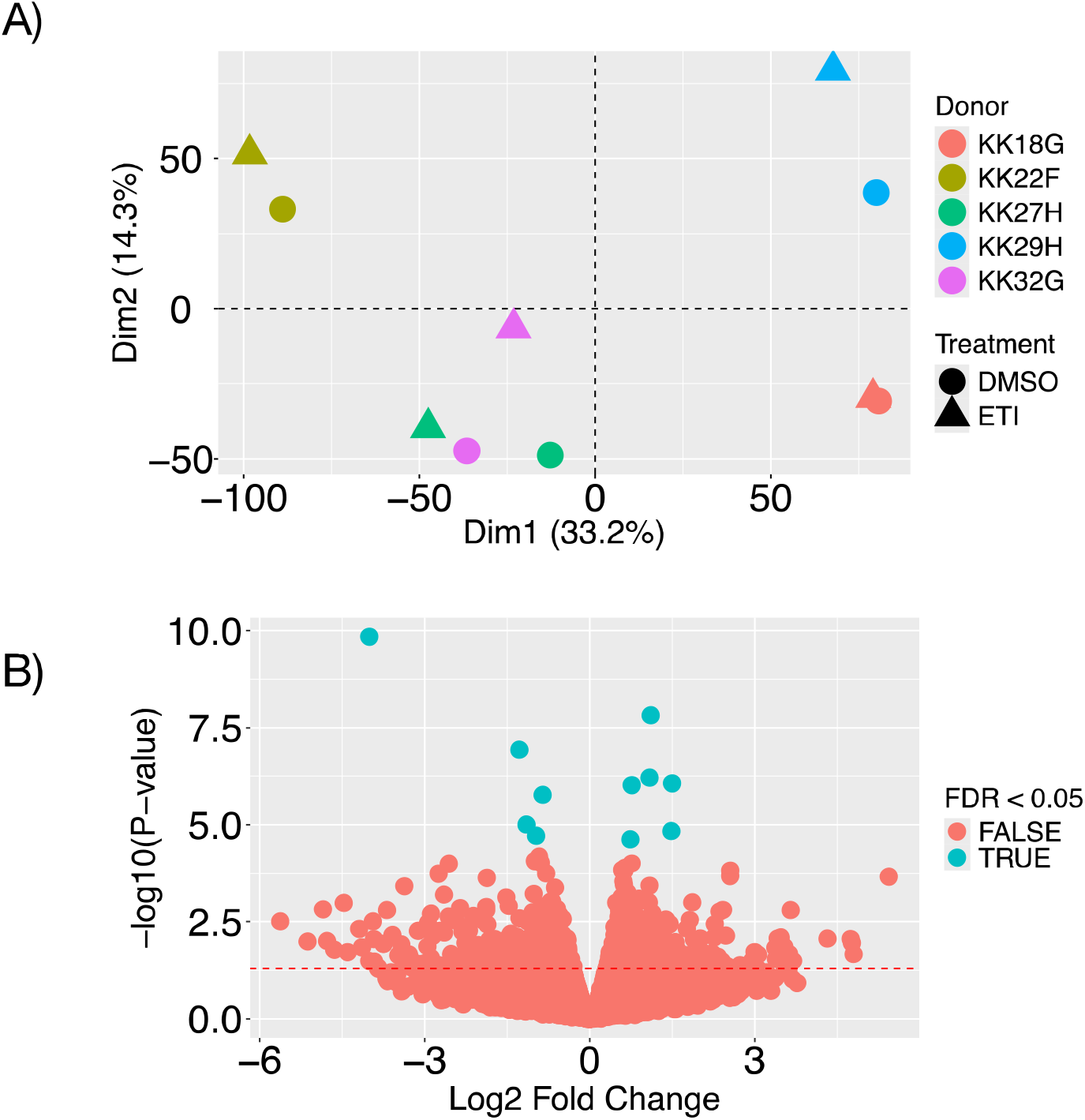
A) Principal component analysis of airway epithelial cells from 5 CF donors exposed to ETI or vehicle (DMSO) for 48 h. Numbers in parentheses indicate the percentage of variability explained by a given principal component (Dim=Dimension). B) Volcano plot of gene expression. 11 genes significantly affected by ETI (FDR <0.05) are indicated in turquoise. *CFTR* gene expression was not significantly affected by ETI (log2FC = -0.27; P = .44).

11 of the 15,283 genes identified with at least 10 counts in any given sample and at least 15 counts across all 10 samples were differentially expressed (FDR < .05) in ETI exposed CF AEC compared to vehicle control as shown in Table 1 and graphically in Figure 1B. It is important to note that others have reported that CFTR modulators like ivacaftor and lumacaftor also have very modest effects on gene expression (38).

**Table 1.**
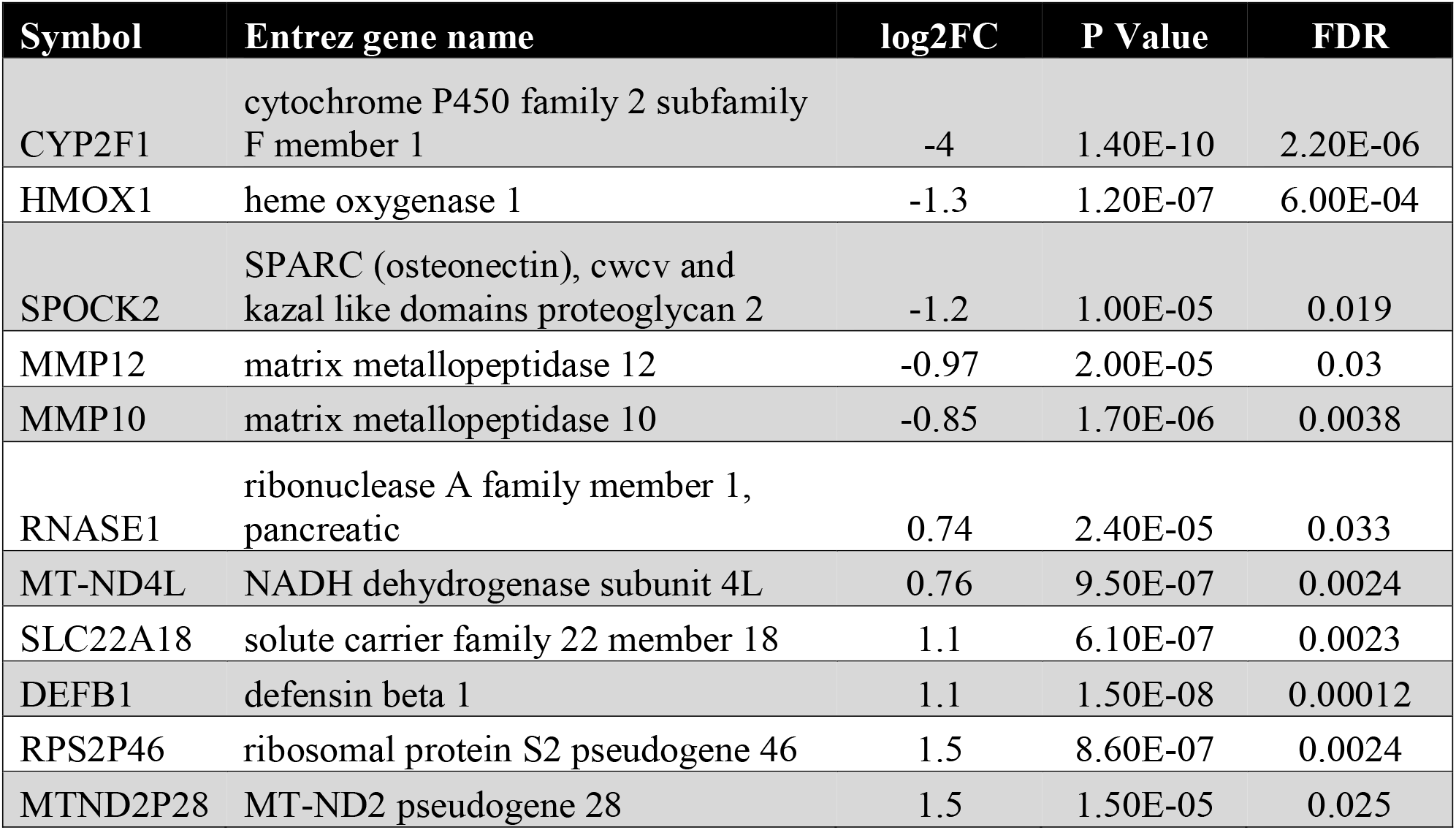
Eleven genes significantly changed by ETI (FDR < .05) ordered by increasing fold change.

Nonetheless, several of the genes that were differentially expressed (FDR < .05) are intriguing in the context of the CF airway. For example, *DEFB1*, which was increased by ETI, codes for human beta defensin 1, an antimicrobial peptide produced by airway epithelial cells (39). *DEFB1* expression is activated in the response to airway pathogens (41) including *Pseudomonas aeruginosa* (42). Reduced pH in the CF airway surface liquid reduces DEFB1 activity (40). However, when modulators like ETI restore pH levels to near normal levels (41) they likely restore DEFB1 activity. Moreover, increased *DEFB1* gene expression coupled with elevated hBD-1 (the protein encoded by *DEFB1*) protein is likely to reduce pathogen abundance and contribute to ETI’s remarkable success (6–9, 41).

Matrix metallopeptidases have a well-established role in CF lung disease since they degrade connective tissue and lead to irreversible lung damage (42) making them attractive targets for drug development (43). Therefore, the observed decrease in the expression of the matrix metallopeptidase genes *MMP10* and *MMP12* by ETI, suggest that ETI may slow lung damage in CF.

Finally, ETI treatment reduced the expression of the stress response gene heme oxygenase (*HMOX1*). Heme oxygenase breaks heme down and releases carbon monoxide (CO), iron and biliverdin (44). These byproducts promote anti-inflammatory signaling in the CF airway, as reviewed by DiPietro et al. (45) and CO is bactericidal against *Pseudomonas aeruginosa* (46). Indeed, *HMOX1* is induced in response to Pseudomonas, and is a known modifier gene of CF expression (47). Notably, expression of *HMOX1* is reduced in CF (48). Therefore, decreased *HMOX1* gene expression in CF airway epithelial cells following ETI suggests that ETI might compromise an anti-bacterial and anti-inflammatory state that is already deficient in CF.

### qPCR confirmation of ETI-induced changes in genes observed by RNA-seq

qPCR was performed to validate significant changes in gene expression observed using RNA-seq. Reassuringly, *DEFB1, MMP10, MMP12*, and *HMOX1* which were differentially expressed as determined by RNA-seq (FDR < .05), were also differentially expressed as assessed by qPCR (Figure 2). *IL1B* and *TNF*, two highly relevant immune genes in CF, were not significantly differentially expressed as determined by both RNA-seq and qPCR.

**Figure 2.**
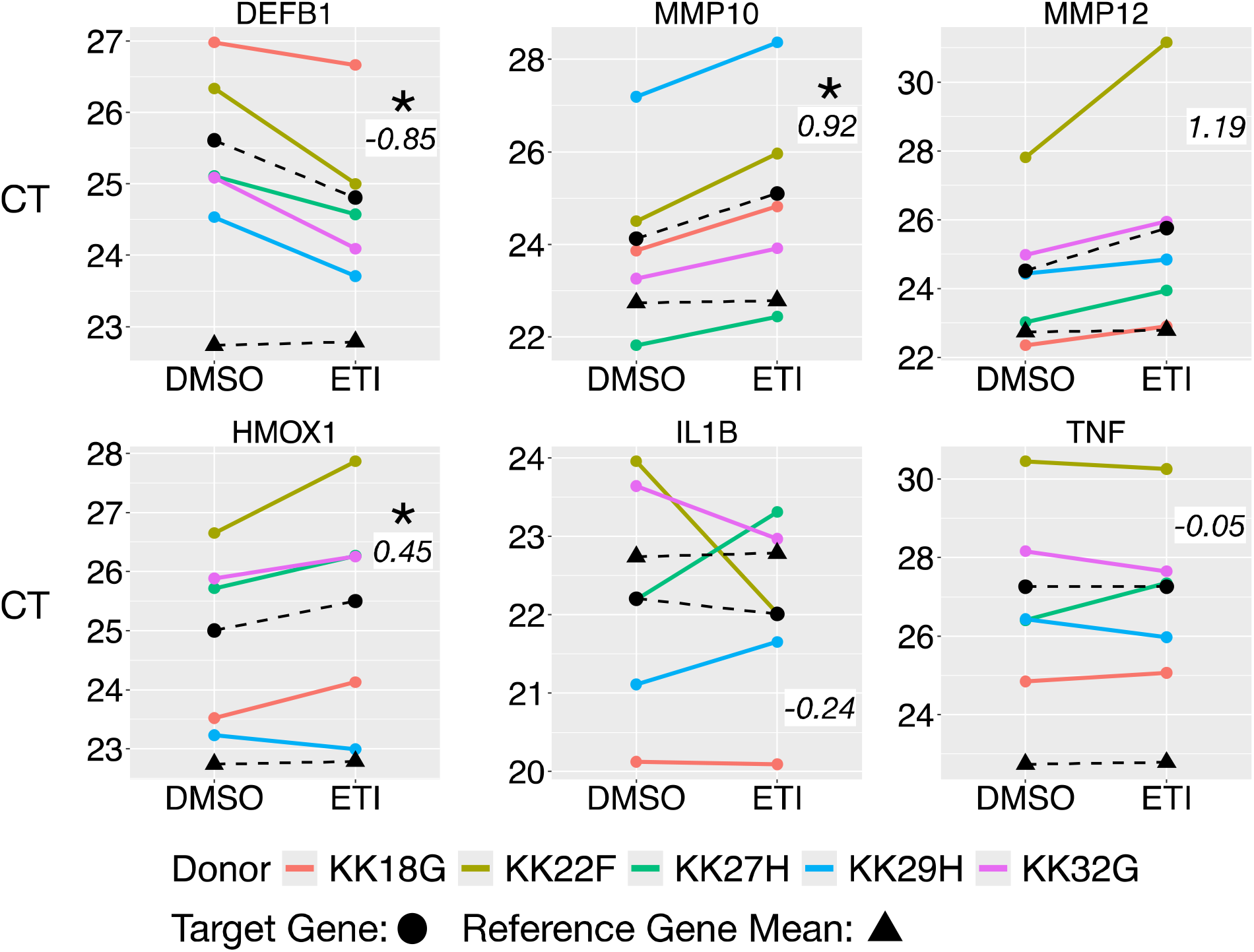
qPCR measurements of *DEFB1, MMP10, MMP12, HMOX1, IL1B, TNF*, and reference genes. Cycle threshold (CT) mean values of each target gene (black circles connected by a dashed line) and reference gene mean values (black triangles connected by a dashed line) are presented. CT values of the gene of interest are shown for each donor (colors). The figure shows that the average of the three reference genes (*HSP90AB, GAPDH, HPRT1)* was neither induced nor repressed in response to ETI: the slope of dashed lines connecting the mean of reference genes in the two conditions is close to zero. *DEFB1, MMP10, MMP12, HMOX1* were significantly differentially expressed in response to ETI using a mixed effect linear model of the ETI effect on each target gene accounting for the mean of the reference genes with donor modeled as a random effect. *DEFB1*: P = .02, *MMP10*: P = .03, *MMP12*: P = .09, *HMOX1*: P = .02, IL1B: P = .79, TNF: P = .99. Delta-delta CT values for each target shown in italics in each panel. Asterisks (*) indicate P < .05. A positive ΔΔCT value of 1 corresponds to a twofold (200%) increase in abundance in the gene of interest.

### Effect of ETI on hBD-1, G-CSF, IL-6, IL-8 and MCP1 secretion by primary CF AEC

High throughput measurement of gene expression readily captures the expression of most genes, but changes in protein expression are more likely to predict changes in phenotype and often gene expression does not correlate with protein expression. For this reason, we used ELISA to assess protein responses to ETI in the same samples that were assessed by RNA-seq and qPCR. Samples were interrogated for several proteins relevant to CF, including hBD-1, G-CSF, IL-6, IL-8 and MCP1. As shown in Figure 3, changes in beta defensin 1 (hBD-1, the protein coded by *DEFB1*) in apical supernatants of CF airway epithelial cells exposed to ETI increased but the increase was not significant (Table 2). It is important to note that some donors appeared to respond to ETI, similar to the variable response of pwCF exposed to ETI (6, 15, 17, 49). For example, ETI increased hBD-1 by 1.12-fold (log_2_) in donor KK29H (turquoise color), a >200% increase above vehicle control. On the other hand, ETI exposure had a much smaller impact on hBD-1 secretion in donors like KK32G (pink color) where hBD-1 levels were at or below baseline levels.

**Table 2.**
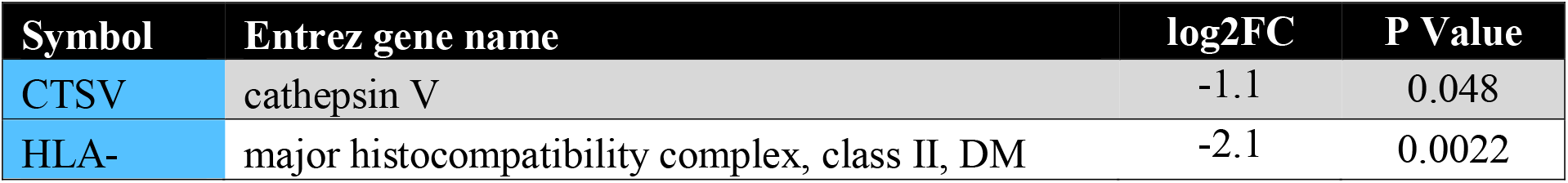
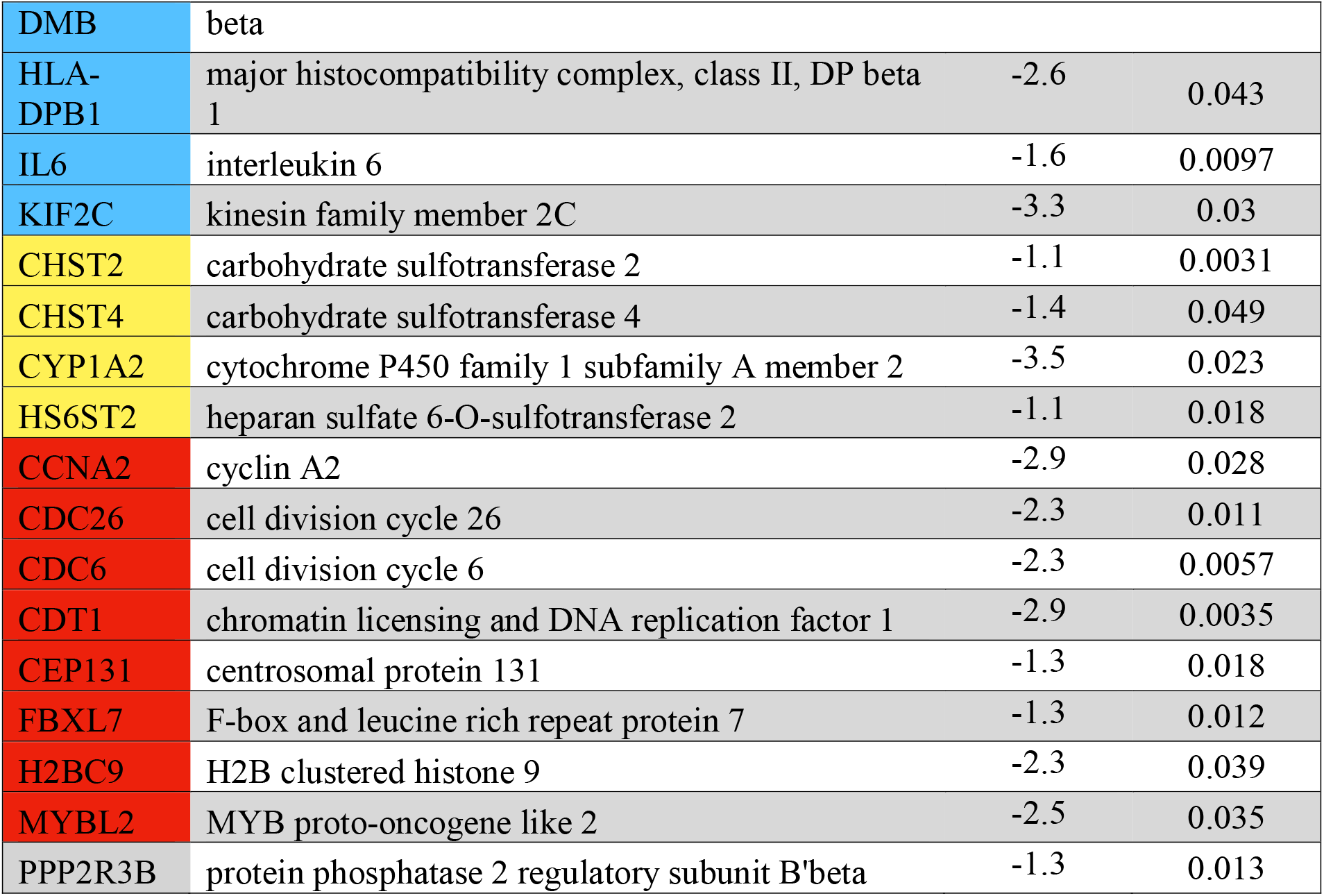
ETI effect on expression of genes shown in Figure 4. Colors indicate the gene pathway to which each gene belongs: blue: Inflammation, yellow: Drug Metabolism, red: Proliferation, Gray: Multiple.

| Symbol | Entrez gene name | log2FC | P Value |
| --- | --- | --- | --- |
| CTSV | cathepsin V | -1.1 | 0.048 |
| HLA- | major histocompatibility complex, class II, DM | -2.1 | 0.0022 |
| DMB | beta |  |  |
| HLA-DPB1 | major histocompatibility complex, class II, DP beta 1 | -2.6 | 0.043 |
| IL6 | interleukin 6 | -1.6 | 0.0097 |
| KIF2C | kinesin family member 2C | -3.3 | 0.03 |
| CHST2 | carbohydrate sulfotransferase 2 | -1.1 | 0.0031 |
| CHST4 | carbohydrate sulfotransferase 4 | -1.4 | 0.049 |
| CYP1A2 | cytochrome P450 family 1 subfamily A member 2 | -3.5 | 0.023 |
| HS6ST2 | heparan sulfate 6-O-sulfotransferase 2 | -1.1 | 0.018 |
| CCNA2 | cyclin A2 | -2.9 | 0.028 |
| CDC26 | cell division cycle 26 | -2.3 | 0.011 |
| CDC6 | cell division cycle 6 | -2.3 | 0.0057 |
| CDT1 | chromatin licensing and DNA replication factor 1 | -2.9 | 0.0035 |
| CEP131 | centrosomal protein 131 | -1.3 | 0.018 |
| FBXL7 | F-box and leucine rich repeat protein 7 | -1.3 | 0.012 |
| H2BC9 | H2B clustered histone 9 | -2.3 | 0.039 |
| MYBL2 | MYB proto-oncogene like 2 | -2.5 | 0.035 |
| PPP2R3B | protein phosphatase 2 regulatory subunit B'beta | -1.3 | 0.013 |

**Figure 3.**
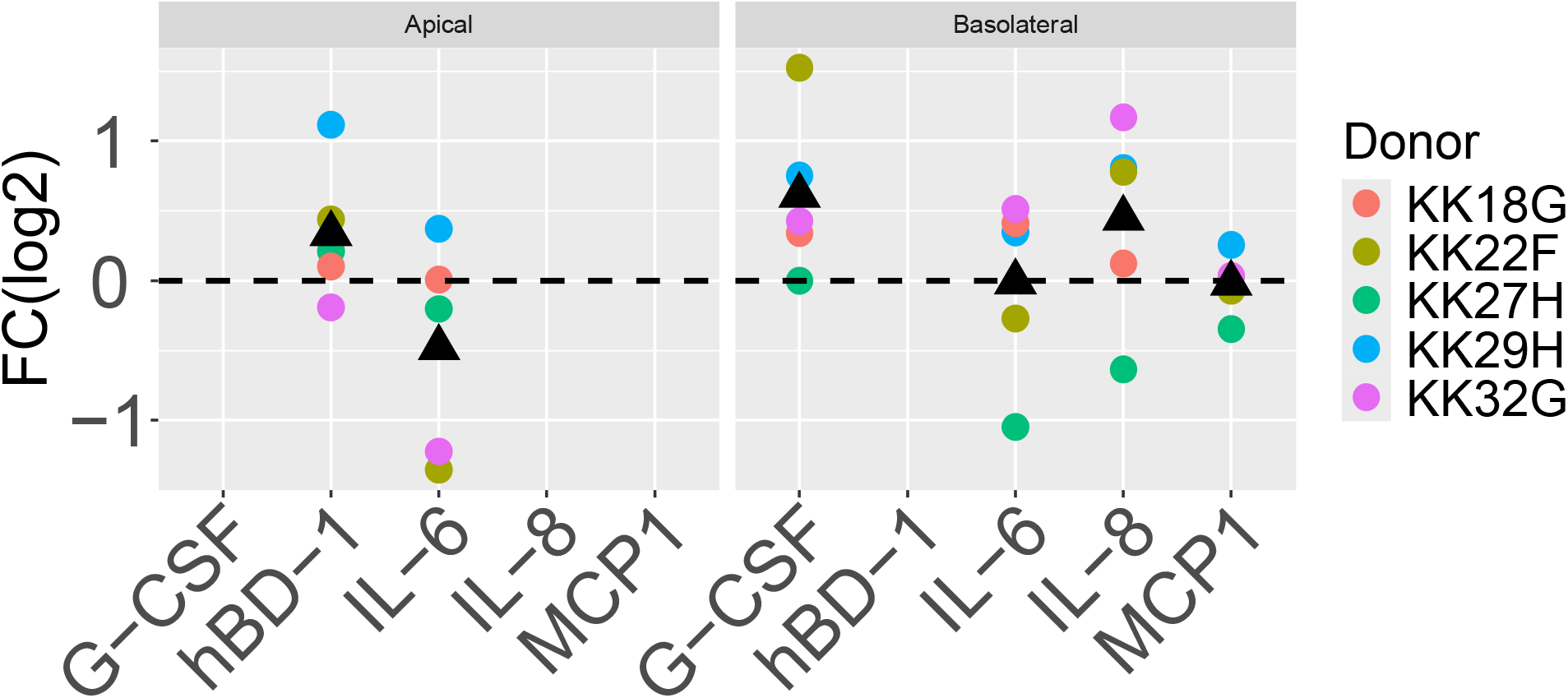
Log base 2 fold change [FC(log2)] responses of beta defensin 1 (hBD-1) and cytokines measured by ELISA to ETI are shown for each analyte detected in the apical fluid (left) or the basolateral solution (right) in each donor (colors) as well as the mean of all donors (black triangles). Significance was assessed by one tailed t tests in which the alternative hypothesis was selected based on prior RNA-seq measurements. Significance is annotated for tests with P < 0.2. There was not enough fluid on the apical side of AEC in air liquid interface (ALI) culture to measure more than hBD-1 and G-CSF.

Donor to donor variability was also observed in the secretion of G-CSF, IL-6, IL-8 and MCP1 in the basolateral media in response to ETI (Figure 3). Only G-CSF, which enhances neutrophil mediated immunity (50) was significantly increased by ETI (P=0.04). G-CSF was detected in basolateral supernatants (right panel of Figure 3) and its response to ETI was concordant with gene expression responses, with an average increase in G-CSF of 150% (P = .04). Donor KK22F (olive) showed the greatest increase in basolateral G-CSF, achieving a 288% increase relative to control.

### Gene pathway analysis by Ingenuity Pathway analysis

The preceding sections focused on the behavior of individual genes, that were significantly altered by ETI (FDR < .05), and the proteins the genes encode. However, groups of genes (e.g., genes belonging to the same biological pathway), work together to maintain homeostasis and respond to pathogens and drugs. We therefore hypothesized that exposing AEC to ETI might target genes in specific biological pathways more than expected by chance, especially if we relax our definition of significance to include any gene with a nominal P value less than .05 and an absolute log2 fold change greater than 1, as is customary in pathway analysis(51, 52). This hypothesis is readily tractable using over-representation analysis in Ingenuity Pathway Analysis (IPA: QIAGEN Inc., https://digitalinsights.qiagen.com/IPA) by Fisher’s exact test. IPA identifies gene pathways in which an experimental condition (e.g., ETI) affects expression of multiple genes in canonical gene pathways. Achieving significance in overrepresentation on a particular path does not require that every gene in the pathway is significantly changed by the treatment, but rather that a significantly higher proportion of differentially expressed genes belong to the pathway than would be predicted by chance.

The results of IPA analysis are summarized in Figure 4, which shows the genes (columns) that participated in the identification of IPA canonical pathways (rows) that had a significantly higher proportion of differentially expressed genes than predicted by chance. Using a less restrictive definition of significance (nominal P < .05 and absolute log2 fold change greater than 1) 288 genes were used as input for overrepresentation analysis in IPA. 178 of these genes were repressed and 103 were induced in response to ETI. Figure 4 only includes pathways that were systematically biased toward induction or repression, i.e., they had an IPA activation z score greater than 1 or less than -1.

**Figure 4.**
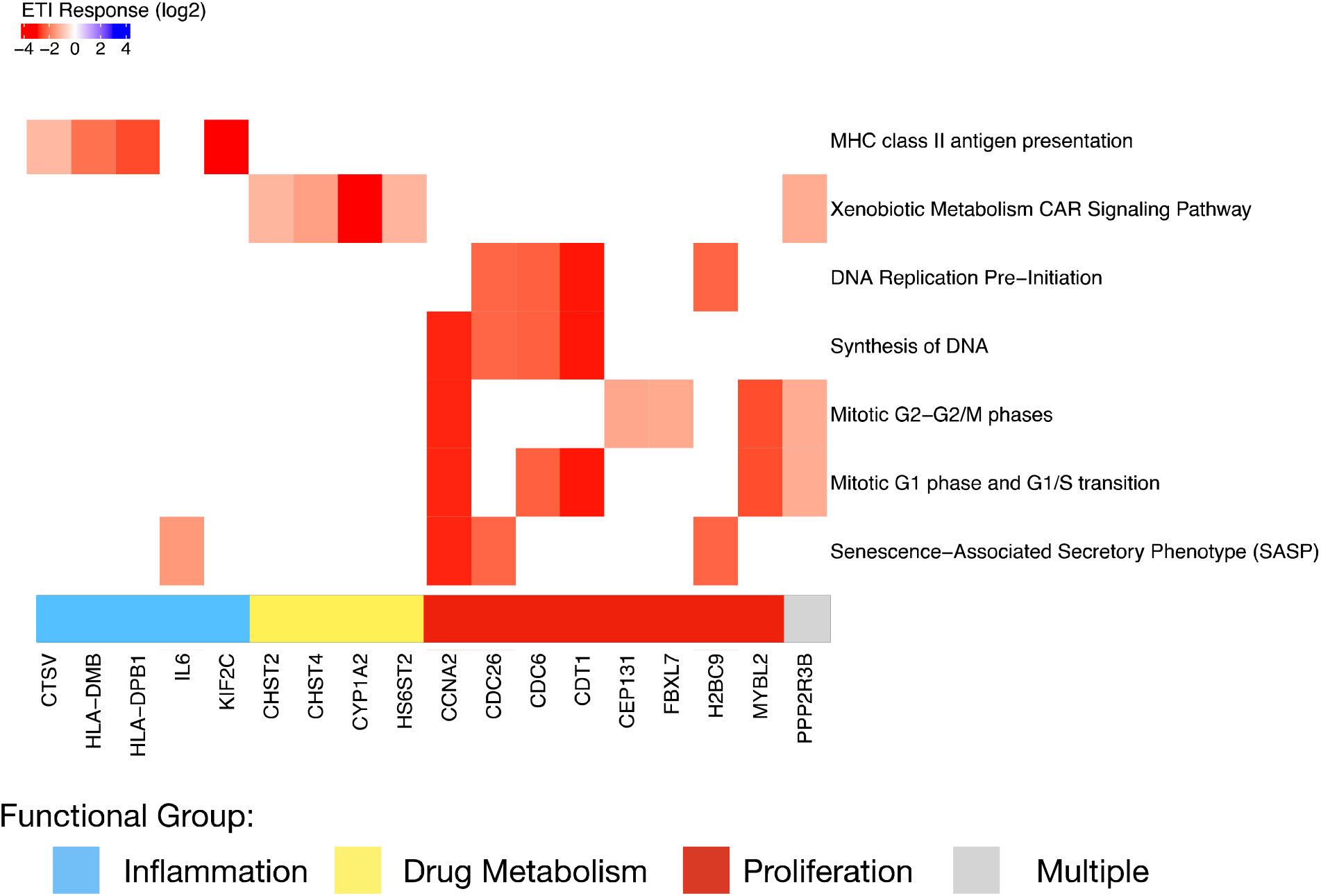
IPA canonical pathways that are relevant to airway epithelial cells. All pathways in the figure are repressed (absolute z-score > 1), and significantly enriched (P < .05, Fisher’s exact test). Pathway names are shown on the right and the gene symbols of genes mapping to each pathway that were differentially expressed are shown on the x axis. Blanks (white color) indicate that a gene is not on a pathway. For example, cathepsin V (CTSV, far left) was significantly repressed by ETI and is a member of the MHC Class II antigen presentation pathway, but not other pathways in this figure.

Figure 4 illustrates three general features of the analysis. First, all of the data in Figure 4 are colored red, meaning that differentially expressed genes mapping to targeted pathways were all repressed. Deeper red colors indicate greater repression in response to ETI. Second, a relatively small number of differentially expressed genes is responsible for pathway enrichment. For example, MHC Class II antigen presentation in the top row of Figure 4 was significantly enriched based on differential expression of four genes *CSTV, HLA-DMB, HLA-DBP1*, and *KIF2C*. Third, the same differentially expressed genes contribute to the significant enrichment of several pathways that are biologically related to each other, which are color coded at the bottom of Figure 4 and described in Table 2.

### Decreased inflammatory signaling

ETI reduced inflammatory signaling by CF AEC as evidenced by significantly reduced expression (log2 fold change < 1, P < .05) of *CSTV, HLA-DMB, HLA-DBP1*, and *IL6*. The IL-6 cytokine was repressed in the apical media collected from airway epithelial cells exposed to ETI, but the change was not significant (Figure 3 and Table 2).

### Decreased xenobiotic response

ETI reduced *CHST2, CHST4, CYP1A2, HS6ST2*, and *PPP2R3B* on the Xenobiotic Metabolism CAR Signaling Pathway as shown in Figure 4. Airway epithelial cells efficiently metabolize drugs (53) and it is possible that repressed xenobiotic metabolism could alter drug-drug interactions in the airway. Sufficient concentrations of drugs can trigger phase I and phase II detoxification processes in human airway epithelial cells (54). Systematic repression of *CHST2, CHST4, CYP1A2, HS6ST2*, and *PPP2R3B* therefore argues against the hypothesis that ETI places a significant toxic burden on airway epithelial cells.

### Decreased cellular proliferation

ETI decreased cell proliferation genes associated with DNA replication, synthesis, and mitotic phase transition (*CCNA, CDC26, CDC6, CDT1, CEP131, FBXL7, H2BC9, H2BC9, and MYBL2*) suggesting that ETI exposure decreases proliferation of CF AEC.

## Discussion

Taken together our analysis of primary human CF AEC reveals that ETI may improve patient outcomes in part by enhancing antibacterial activity, reducing lung damage, and suppressing proinflammatory signaling (Figure 4 and Table 2). Collectively, the data suggest that the ETI induced reduction in lung infections in pwCF are related in part to drug induced increases in *DEFB1*, and that ETI may reduce lung damage by reducing *MMP10* and *MMP12* gene expression, which is predicted to reduce matrix metalloprotease activity. Moreover, pathway analysis also identified several genes responsible for the ETI induced reduction in inflammation observed in people with CF. Our data are consistent with literature reports that ETI reduces bacterial burden and inflammation in the CF airway (55), improves lung function, and reduces exacerbations and hospitalizations with few side effects (7, 9, 41, 55).

This study identifies mechanisms through which ETI is likely to improve antibacterial function, reduce lung damage, and reduce proinflammatory signaling. Specifically, we found that ETI increased gene expression of DEFB1 and reduced the expression of genes mediating inflammation and lung damage. Beneficial effects of other CFTR modulators have been previously reported (11), and that subjects with the largest gene expression responses to ivacaftor experience the greatest clinical benefit (12), and that the combination of ivacaftor and lumacaftor decreased the expression of cell-death genes, *MMP9* and *SOCS3* (13). We have shown previously that exposing CF monocyte derived macrophages to lumacaftor and ivacaftor reduced transferrin receptor 1 expression, corrects dysfunctional iron transport gene expression observed in CF (56), and that lumacaftor restores the ability of CF macrophages to phagocytose and kill Pseudomonas (57).

Two recent studies have examined the effect of ETI on the transcriptional response of nasal swabs obtained from pwCF (15, 17). Loske et al reported that ETI improves innate immunity and suppresses immune cell inflammatory responses in children with CF (15). Yue et al. identified inverse correlations between inflammatory gene expression (e.g., *TLR4* and *IL10)* and FEV1, but they did not find that *DEFB1, MMP10, MMP12, HMOX1* (genes we identified as responsive to ETI) were ideal biomarkers to predict clinical outcomes in pwCF (17).

As noted above in Figure 1A, most of the variation in the data is attributable to donor differences rather than ETI exposure. This is not surprising given that pwCF respond to ETI in varying degrees, some have robust responses and a some have small or negligible responses (6, 7, 9, 41, 55). Moreover, many studies that have examined the effect of other modulators of CFTR and have shown very small or no effects on gene expression (12, 13, 38). It should be noted that predictions based primarily on *in vitro* gene expression, and ELISA analysis of cytokine secretion by CF AEC cells needs to be validated by clinical measurements of cytokines in bronchoalveolar fluid (BALF) and/or sputum (if available) in pwCF. Several studies have shown in pwCF that ETI reduces *Pseudomonas* abundance and systemic inflammation (7, 58–62).

However, the effect of ETI on gene expression in lung tissue of pwCF has, to the best of our knowledge, not been reported.

### DEFB1, ETI and lung infection

Antimicrobial peptides like beta defensins are effective against CF pathogens (63), are secreted by airway epithelial cells, are present in CF bronchoalveolar lavage fluid (39), and predictive of lung disease severity (64). Genetic polymorphisms in *DEFB1* have been proposed to explain differences in colonization by *P. aeruginosa* in pwCF (65). Beta defensin 1 is pH sensitive (40), thus enhancement of the ability of CFTR to secrete Cl^-^ and HCO_3_^-^ and increase airway surface fluid pH as well as an increase in DEFB1 expression as a results of ETI exposure is predicted to increase the ability of hBD-1 to kill bacteria.

### Decreased MMP10 and MMP12 by ETI in the CF lung is predicted to reduce lung damage

MMPs are expressed in and secreted by airway epithelial cells (66) and play a key role in extracellular matrix remodeling by degrading extracellular matrix components. A pathogenic role for matrix metalloproteinase 12 (MMP12) in lung disease is supported by multiple lines of evidence (67). MMPs stimulate pro-inflammatory cytokines including TNFα (68) and exacerbate chronic lung infection and inflammation by promoting tissue damage and fibrosis (69). MMP12 is upregulated in βENaC-Tg mice and contributes to early lung damage (67). MMPs also cleave inactive proforms of cytokines, such as pro-IL-1β, into their active forms, thereby potentiating inflammation (70). *P. aeruginosa* induced increases in IL-6, TNFα, and IL-8 upregulate MMP-12 and thereby promote a proteolytic environment that facilitates lung destruction (69). Thus, a reduction in MMP10 and MMP12 by ETI in the CF lung is predicted to reduce lung damage in CF.

Matrix metalloprotease MMP10 is also upregulated in AEC exposed to *P. aeruginosa* and by airway pathogens in general (71–73) and in a mouse model of *P. aeruginosa* lung infection (71). *Mmp10*^−/−^ mice are more susceptible to pneumonia than wild-type mice (71). MMP10 facilitates the activation of macrophages from proinflammatory (M1-like) cells into immunosuppressive (M2-like) cells (74). ETI-mediated reduction of MMP10 is therefore also predicted to benefit people with CF.

### Decreased inflammatory signaling induced by ETI

ETI reduced the expression of a cluster of genes with shared pro-inflammatory function (*CSTV, HLA-DMB, HLA-DBP1, IL6*) (Figure 4), consistent with other studies demonstrating that ETI reduces inflammation in pwCF (75).

### The current study has several strengths

To our knowledge, this is the first study to comprehensively examine the effect of ETI on primary CF AEC in air liquid interface culture. Our results are in general agreement with other studies on blood and nasal swabs that ETI reduces inflammation. In addition, we identified several genes including *DEFB1, MMP10, MMP12, HMOX1* and several proinflammatory immune pathways that are down regulated by ETI. Using qPCR we confirmed the differential expression of these genes identified by RNA-seq. Notably, as suggested previously, we identified several reference genes that when averaged were not affected by ETI, and we presented the data on the CT values of reference genes as well as our genes of interest to demonstrate that the changes in ΔΔCT were due to changes in genes of interest as opposed to fluctuations in reference genes (18). Moreover, our studies were conducted on primary cultures of CF AEC in air liquid interface culture from five donors. Given the variability of the response of pwCF to ETI, we suggest that our studies on primary cells from five donors are likely to be more representative of the disease than studies that interrogate cell lines that have been obtained from a single donor, some of which were obtained from tumors or have been genetically manipulated.

### We also acknowledge limitations

First, in this study we studied five donors of CF AEC cells. It is possible that studies using more donors would uncover small changes in gene and/or cytokine expression that did not achieve statistical significance in this study. However, a power analysis of our data reveal that ∼17 donors would be needed to identify small changes (P = .05, with 80% power) in several genes and IL-6 and other cytokines that were affected by ETI but did not reach statistical significance (Figure 3). Second, as with all studies on primary AEC obtained from pwCF, and on studies on pwCF, previous exposure to CFTR modulators and/or other medications, as well as environment factors and culture conditions may affect their response to ETI and thereby account for some of the observations obtained in this study. Thus, it is possible that studies on another cohort of AEC may identify additional ETI effects. However, given that our studies on ETI are in general agreement (e.g., ETI reduces inflammation) with studies on blood and nasal samples from pwCF, we do not believe this is not a significant limitation.

### Conclusions

Our study of gene expression responses by CF AEC exposed to ETI suggest that in addition to improving CFTR channel function, ETI is likely to increase resistance to bacterial infection, and thereby account for the modest reduction in bacteria by increasing hBD-1, and reducing lung damage by suppressing MMP10, and reducing inflammation by repressing proinflammatory cytokine secretion by AEC cells. Furthermore, the reduction in proinflammatory cytokines is predicted to reduce inflammation by decreasing the recruitment of immune cells, which are known to induce lung damage, into the lungs.

## Acknowledgements

This work was supported by funding from the Cystic Fibrosis Foundation (STANTO19G0, STANTO20P0, and STANTO19R0), from the National Institutes of Health (P30DK117469 and R01HL151385) and the Flatley Foundation.

We are grateful to the Genomic Data Sciences core of the Center for Quantitative Biology at Dartmouth for preprocessing and initial analysis of bulk RNA-seq data, which receives support from 5P20GM130454-03 and the Dartmouth Cancer Center through NCI Cancer Center Support Grant 5P30 CA023108-43.

## References

1. Stanton BA. Effects of Pseudomonas aeruginosa on CFTR chloride secretion and the host immune response. American Journal of Physiology-Cell Physiology 312: C357–C366, 2017. doi: 10.1152/ajpcell.00373.2016.

2. Grasemann H, Ratjen F. Cystic Fibrosis. New England Journal of Medicine 389: 1693–1707, 2023. doi: 10.1056/NEJMra2216474.

3. Tice JA, Kuntz KM, Wherry K, Seidner M, Rind DM, Pearson SD. The effectiveness and value of novel treatments for cystic fibrosis. J Manag Care Spec Pharm 27: 10.18553/jmcp.2021.27.2.276, 2021. doi: 10.18553/jmcp.2021.27.2.276.

4. Connett G. Lumacaftor-ivacaftor in the treatment of cystic fibrosis: design, development and place in therapy. Drug Design, Development and Therapy 13: 2405–2412, 2019. doi: 10.2147/DDDT.S153719.

5. Taylor-Cousar Jennifer L., Munck Anne, McKone Edward F., van der Ent Cornelis K., Moeller Alexander, Simard Christopher, Wang Linda T., Ingenito Edward P., McKee Charlotte, Lu Yimeng, Lekstrom-Himes Julie, Elborn J. Stuart. Tezacaftor– Ivacaftor in Patients with Cystic Fibrosis Homozygous for Phe508del. New England Journal of Medicine 377: 2013–2023, 2017. doi: 10.1056/NEJMoa1709846.

6. Middleton Peter G., Mall Marcus A., Dřevínek Pavel, Lands Larry C., McKone Edward F., Polineni Deepika, Ramsey Bonnie W., Taylor-Cousar Jennifer L., Tullis Elizabeth, Vermeulen François, Marigowda Gautham, McKee Charlotte M., Moskowitz Samuel M., Nair Nitin, Savage Jessica, Simard Christopher, Tian Simon, Waltz David, Xuan Fengjuan, Rowe Steven M., Jain Raksha. Elexacaftor–Tezacaftor– Ivacaftor for Cystic Fibrosis with a Single Phe508del Allele. New England Journal of Medicine 381: 1809–1819, 2019. doi: 10.1056/NEJMoa1908639.

7. Bacalhau M, Camargo M, Magalhães-Ghiotto GAV, Drumond S, Castelletti CHM, Lopes-Pacheco M. Elexacaftor-Tezacaftor-Ivacaftor: A Life-Changing Triple Combination of CFTR Modulator Drugs for Cystic Fibrosis. Pharmaceuticals (Basel) 16: 410, 2023. doi: 10.3390/ph16030410.

8. Heijerman HGM, McKone EF, Downey DG, Van Braeckel E, Rowe SM, Tullis E, Mall MA, Welter JJ, Ramsey BW, McKee CM, Marigowda G, Moskowitz SM, Waltz D, Sosnay PR, Simard C, Ahluwalia N, Xuan F, Zhang Y, Taylor-Cousar JL, McCoy KS, VX17-445-103 Trial Group. Efficacy and safety of the elexacaftor plus tezacaftor plus ivacaftor combination regimen in people with cystic fibrosis homozygous for the F508del mutation: a double-blind, randomised, phase 3 trial. Lancet 394: 1940–1948, 2019. doi: 10.1016/S0140-6736(19)32597-8.

9. Zemanick ET, Taylor-Cousar JL, Davies J, Gibson RL, Mall MA, McKone EF, McNally P, Ramsey BW, Rayment JH, Rowe SM, Tullis E, Ahluwalia N, Chu C, Ho T, Moskowitz SM, Noel S, Tian S, Waltz D, Weinstock TG, Xuan F, Wainwright CE, McColley SA. A Phase 3 Open-Label Study of Elexacaftor/Tezacaftor/Ivacaftor in Children 6 through 11 Years of Age with Cystic Fibrosis and at Least One F508del Allele. Am J Respir Crit Care Med 203: 1522–1532, [date unknown]. doi: 10.1164/rccm.202102-0509OC.

10. Becq F, Mirval S, Carrez T, Lévêque M, Billet A, Coraux C, Sage E, Cantereau A. The rescue of F508del-CFTR by elexacaftor/tezacaftor/ivacaftor (Trikafta) in human airway epithelial cells is underestimated due to the presence of ivacaftor. European Respiratory Journal 59, 2022. doi: 10.1183/13993003.00671-2021.

11. Hisert KB, Birkland TP, Schoenfelt KQ, Long ME, Grogan B, Carter S, Liles WC, McKone EF, Becker L, Manicone AM, Gharib SA. CFTR Modulator Therapy Enhances Peripheral Blood Monocyte Contributions to Immune Responses in People With Cystic Fibrosis. Front Pharmacol 11: 1219, 2020. doi: 10.3389/fphar.2020.01219.

12. Sun T, Sun Z, Jiang Y, Ferguson AA, Pilewski JM, Kolls JK, Chen W, Chen K. Transcriptomic Responses to Ivacaftor and Prediction of Ivacaftor Clinical Responsiveness. Am J Respir Cell Mol Biol 61: 643–652, 2019. doi: 10.1165/rcmb.2019-0032OC.

13. Kopp BT, Fitch J, Jaramillo L, Shrestha CL, Robledo-Avila F, Zhang S, Palacios S, Woodley F, Hayes D, Partida-Sanchez S, Ramilo O, White P, Mejias A. Whole-blood transcriptomic responses to lumacaftor/ivacaftor therapy in cystic fibrosis. Journal of Cystic Fibrosis 19: 245–254, 2020. doi: 10.1016/j.jcf.2019.08.021.

14. Yang Q, Soltis AR, Sukumar G, Zhang X, Caohuy H, Freedy J, Dalgard CL, Wilkerson MD, Pollard HB, Pollard BS. Gene therapy-emulating small molecule treatments in cystic fibrosis airway epithelial cells and patients. Respiratory Research 20: 290, 2019. doi: 10.1186/s12931-019-1214-8.

15. Loske J, Völler M, Lukassen S, Stahl M, Thürmann L, Seegebarth A, Röhmel J, Wisniewski S, Messingschlager M, Lorenz S, Klages S, Eils R, Lehmann I, Mall MA, Graeber SY, Trump S. Pharmacological Improvement of Cystic Fibrosis Transmembrane Conductance Regulator Function Rescues Airway Epithelial Homeostasis and Host Defense in Children with Cystic Fibrosis. Am J Respir Crit Care Med 209: 1338–1350, 2024. doi: 10.1164/rccm.202310-1836OC.

16. Idris T, Bachmann M, Bacchetta M, Wehrle-Haller B, Chanson M, Badaoui M. Akt-driven TGF-β and DKK1 Secretion Impairs F508del CF Airway Epithelium Polarity. Am J Respir Cell Mol Biol Epub ahead of print, 2024. doi: 10.1165/rcmb.2023-0408OC.

17. Yue M, Weiner DJ, Gaietto KM, Rosser FJ, Qoyawayma CM, Manni ML, Myerburg MM, Pilewski JM, Celedón JC, Chen W, Forno E. Nasal Epithelium Transcriptomics Predict Clinical Response to Elexacaftor/Tezacaftor/Ivacaftor. American Journal of Respiratory Cell and Molecular Biology Epub ahead of print, 2024. doi: 10.1165/rcmb.2024-0103OC.

18. Hampton TH, Koeppen K, Bashor L, Stanton BA. Selection of reference genes for quantitative PCR: identifying reference genes for airway epithelial cells exposed to Pseudomonas aeruginosa. Am J Physiol Lung Cell Mol Physiol 319: L256–L265, 2020. doi: 10.1152/ajplung.00158.2020.

19. Barnaby R, Koeppen K, Stanton BA. Cyclodextrins reduce the ability of Pseudomonas aeruginosa outer-membrane vesicles to reduce CFTR Cl-secretion. Am J Physiol Lung Cell Mol Physiol 316: L206–L215, 2019. doi: 10.1152/ajplung.00316.2018.

20. Stanton BA, Coutermarsh B, Barnaby R, Hogan D. Pseudomonas aeruginosa Reduces VX-809 Stimulated F508del-CFTR Chloride Secretion by Airway Epithelial Cells. PLoS ONE 10, 2015. doi: 10.1371/journal.pone.0127742.

21. Swiatecka-Urban A, Moreau-Marquis S, Maceachran DP, Connolly JP, Stanton CR, Su JR, Barnaby R, O’toole GA, Stanton BA. Pseudomonas aeruginosa inhibits endocytic recycling of CFTR in polarized human airway epithelial cells. Am J Physiol Cell Physiol 290: C862–872, 2006. doi: 10.1152/ajpcell.00108.2005.

22. Martin M. Cutadapt removes adapter sequences from high-throughput sequencing reads. EMBnet.journal 17: 10–12, 2011. doi: 10.14806/ej.17.1.200.

23. Kim D, Paggi JM, Park C, Bennett C, Salzberg SL. Graph-based genome alignment and genotyping with HISAT2 and HISAT-genotype. Nat Biotechnol 37: 907–915, 2019. doi: 10.1038/s41587-019-0201-4.

24. Liao Y, Smyth GK, Shi W. featureCounts: an efficient general purpose program for assigning sequence reads to genomic features. Bioinformatics 30: 923–930, 2014. doi: 10.1093/bioinformatics/btt656.

25. Li H, Handsaker B, Wysoker A, Fennell T, Ruan J, Homer N, Marth G, Abecasis G, Durbin R, 1000 Genome Project Data Processing Subgroup. The Sequence Alignment/Map format and SAMtools. Bioinformatics 25: 2078–2079, 2009. doi: 10.1093/bioinformatics/btp352.

26. Robinson MD, McCarthy DJ, Smyth GK. edgeR: a Bioconductor package for differential expression analysis of digital gene expression data. Bioinformatics 26: 139–140, 2010. doi: 10.1093/bioinformatics/btp616.

27. Oksanen J, Simpson GL, Blanchet FG, Kindt R, Legendre P, Minchin PR, O’Hara RB, Solymos P, Stevens MHH, Szoecs E, Wagner H, Barbour M, Bedward M, Bolker B, Borcard D, Carvalho G, Chirico M, Caceres MD, Durand S, Evangelista HBA, FitzJohn R, Friendly M, Furneaux B, Hannigan G, Hill MO, Lahti L, McGlinn D, Ouellette M-H, Cunha ER, Smith T, Stier A, Braak Cjft, Weedon J. vegan: Community Ecology Package [Online]. 2024. https://cran.r-project.org/web/packages/vegan/index.html [30 May 2024].

28. Kassambara A, Mundt F. factoextra: Extract and Visualize the Results of Multivariate Data Analyses [Online]. 2020. https://CRAN.R-project.org/package=factoextra.

29. Edgar R, Domrachev M, Lash AE. Gene Expression Omnibus: NCBI gene expression and hybridization array data repository. Nucleic Acids Res 30: 207–210, 2002. doi: 10.1093/nar/30.1.207.

30. Chen G, Sun L, Kato T, Okuda K, Martino MB, Abzhanova A, Lin JM, Gilmore RC, Batson BD, O’Neal YK, Volmer AS, Dang H, Deng Y, Randell SH, Button B, Livraghi-Butrico A, Kesimer M, Ribeiro CMP, O’Neal WK, Boucher RC. IL-1β dominates the promucin secretory cytokine profile in cystic fibrosis. J Clin Invest 129: 4433–4450, 2019. doi: 10.1172/JCI125669.

31. Maillé E, Trinh NTN, Privé A, Bilodeau C, Bissonnette É, Grandvaux N, Brochiero E. Regulation of normal and cystic fibrosis airway epithelial repair processes by TNF-α after injury. American Journal of Physiology-Lung Cellular and Molecular Physiology 301: L945–L955, 2011. doi: 10.1152/ajplung.00149.2011.

32. Zhong S. ctrlGene: Assess the Stability of Candidate Housekeeping Genes [Online]. 2019. https://cran.r-project.org/web/packages/ctrlGene/index.html [8 Jun. 2024].

33. Bates D, Maechler M, Bolker [aut B, cre, Walker S, Christensen RHB, Singmann H, Dai B, Scheipl F, Grothendieck G, Green P, Fox J, Bauer A, simulate.formula) PNK (shared copyright on, Tanaka E, Jagan M. lme4: Linear Mixed-Effects Models using “Eigen” and S4 [Online]. 2024. https://cran.r-project.org/web/packages/lme4/index.html [8 Jun. 2024].

34. Kuznetsova A, Brockhoff PB, Christensen RHB. lmerTest Package: Tests in Linear Mixed Effects Models. Journal of Statistical Software 82: 1–26, 2017. doi: 10.18637/jss.v082.i13.

35. Ritz C, Strebig JC. drc: Analysis of Dose-Response Curves [Online]. 2016. https://cran.r-project.org/web/packages/drc/index.html [12 Jun. 2024].

36. R Core Team. R: A language and environment for statistical computing. [Online]. R Foundation for Statistical Computing: 2021. URL https://www.R-project.org/.

37. Wickham H. ggplot2: Elegant Graphics for Data Analysis. Springer-Verlag New York: 2016.

38. De Jong E, Garratt LW, Looi K, Lee AHY, Ling K-M, Smith ML, Falsafi R, Sutanto EN, Hillas J, Iosifidis T, Martinovich KM, Shaw NC, Montgomery ST, Kicic-Starcevich E, Lannigan FJ, Vijayasekaran S, Hancock REW, Stick SM, Kicic A, Arest C. Ivacaftor or lumacaftor/ivacaftor treatment does not alter the core CF airway epithelial gene response to rhinovirus. Journal of Cystic Fibrosis 20: 97–105, 2021. doi: 10.1016/j.jcf.2020.07.004.

39. Singh PK, Jia HP, Wiles K, Hesselberth J, Liu L, Conway B-AD, Greenberg EP, Valore EV, Welsh MJ, Ganz T, Tack BF, McCray PB. Production of β-defensins by human airway epithelia. Proceedings of the National Academy of Sciences 95: 14961–14966, 1998. doi: 10.1073/pnas.95.25.14961.

40. Goldman MJ, Anderson GM, Stolzenberg ED, Kari UP, Zasloff M, Wilson JM. Human β-Defensin-1 Is a Salt-Sensitive Antibiotic in Lung That Is Inactivated in Cystic Fibrosis. Cell 88: 553–560, 1997. doi: 10.1016/S0092-8674(00)81895-4.

41. Nichols DP, Paynter AC, Heltshe SL, Donaldson SH, Frederick CA, Freedman SD, Gelfond D, Hoffman LR, Kelly A, Narkewicz MR, Pittman JE, Ratjen F, Rosenfeld M, Sagel SD, Schwarzenberg SJ, Singh PK, Solomon GM, Stalvey MS, Clancy JP, Kirby S, Van Dalfsen JM, Kloster MH, Rowe SM, PROMISE Study group. Clinical Effectiveness of Elexacaftor/Tezacaftor/Ivacaftor in People with Cystic Fibrosis: A Clinical Trial. Am J Respir Crit Care Med 205: 529–539, 2022. doi: 10.1164/rccm.202108-1986OC.

42. Gaggar A, Hector A, Bratcher PE, Mall MA, Griese M, Hartl D. The role of matrix metalloproteases in cystic fibrosis lung disease. Eur Respir J 38: 721–727, 2011. doi: 10.1183/09031936.00173210.

43. McKelvey MC, Weldon S, McAuley DF, Mall MA, Taggart CC. Targeting Proteases in Cystic Fibrosis Lung Disease. Paradigms, Progress, and Potential. Am J Respir Crit Care Med 201: 141–147, 2020. doi: 10.1164/rccm.201906-1190PP.

44. Kikuchi G, Yoshida T, Noguchi M. Heme oxygenase and heme degradation. Biochemical and Biophysical Research Communications 338: 558–567, 2005. doi: 10.1016/j.bbrc.2005.08.020.

45. Di Pietro C, Öz HH, Murray TS, Bruscia EM. Targeting the Heme Oxygenase 1/Carbon Monoxide Pathway to Resolve Lung Hyper-Inflammation and Restore a Regulated Immune Response in Cystic Fibrosis [Online]. Frontiers in Pharmacology 11, 2020. https://www.frontiersin.org/articles/10.3389/fphar.2020.01059 [20 Sep. 2022].

46. Desmard M, Davidge KS, Bouvet O, Morin D, Roux D, Foresti R, Ricard JD, Denamur E, Poole RK, Montravers P, Morterlini R, Boczkowski J. A carbon monoxide-releasing molecule (CORM-3) exerts bactericidal activity against Pseudomonas aeruginosa and improves survival in an animal model of bacteraemia. The FASEB Journal 23: 1023–1031, 2009. doi: 10.1096/fj.08-122804.

47. Park JE, Yung R, Stefanowicz D, Shumansky K, Akhabir L, Durie PR, Corey M, Zielenski J, Dorfman R, Daley D, Sandford AJ. Cystic fibrosis modifier genes related to Pseudomonas aeruginosa infection. Genes Immun 12: 370–377, 2011. doi: 10.1038/gene.2011.5.

48. Chillappagari S, Garapati V, Mahavadi P, Naehrlich L, Schmeck BT, Schmitz ML, Guenther A. Defective BACH1/HO-1 regulatory circuits in cystic fibrosis bronchial epithelial cells. Journal of Cystic Fibrosis 20: 140–148, 2021. doi: 10.1016/j.jcf.2020.05.006.

49. Alicandro G, Gramegna A, Bellino F, Sciarrabba SC, Lanfranchi C, Contarini M, Retucci M, Daccò V, Blasi F. Heterogeneity in response to Elexacaftor/Tezacaftor/Ivacaftor in people with cystic fibrosis. J Cyst Fibros Epub ahead of print: S1569-1993(24)00057–2, 2024. doi: 10.1016/j.jcf.2024.04.013.

50. Martin KR, Wong HL, Witko-Sarsat V, Wicks IP. G-CSF - A double edge sword in neutrophil mediated immunity. Semin Immunol 54: 101516, 2021. doi: 10.1016/j.smim.2021.101516.

51. Pattison SH, Gibson DS, Johnston E, Peacock S, Rivera K, Tunney MM, Pappin DJ, Elborn JS. Proteomic profile of cystic fibrosis sputum cells in adults chronically infected with Pseudomonas aeruginosa. European Respiratory Journal 50, 2017. doi: 10.1183/13993003.01569-2016.

52. Li Z, Barnaby R, Nymon A, Roche C, Koeppen K, Ashare A, Hogan DA, Gerber SA, Taatjes DJ, Hampton TH, Stanton BA. P. aeruginosa tRNA-fMet halves secreted in outer membrane vesicles suppress lung inflammation in cystic fibrosis. American Journal of Physiology-Lung Cellular and Molecular Physiology 326: L574–L588, 2024. doi: 10.1152/ajplung.00018.2024.

53. Forbes B. Human airway epithelial cell lines for in vitro drug transport and metabolism studies. Pharmaceutical Science & Technology Today 3: 18–27, 2000. doi: 10.1016/S1461-5347(99)00231-X.

54. Boei Jjwa, Vermeulen S, Klein B, Hiemstra PS, Verhoosel RM, Jennen DGJ, Lahoz A, Gmuender H, Vrieling H. Xenobiotic metabolism in differentiated human bronchial epithelial cells. Arch Toxicol 91: 2093–2105, 2017. doi: 10.1007/s00204-016-1868-7.

55. Casey M, Gabillard-Lefort C, McElvaney OF, McElvaney OJ, Carroll T, Heeney RC, Gunaratnam C, Reeves EP, Murphy MP, McElvaney NG. Effect of elexacaftor/tezacaftor/ivacaftor on airway and systemic inflammation in cystic fibrosis. Thorax 78: 835–839, 2023. doi: 10.1136/thorax-2022-219943.

56. Hazlett HF, Hampton TH, Aridgides DS, Armstrong DA, Dessaint JA, Mellinger DL, Nymon AB, Ashare A. Altered iron metabolism in cystic fibrosis macrophages: the impact of CFTR modulators and implications for Pseudomonas aeruginosa survival. Sci Rep 10: 10935, 2020. doi: 10.1038/s41598-020-67729-5.

57. Barnaby R, Koeppen K, Nymon A, Hampton TH, Berwin B, Ashare A, Stanton BA. Lumacaftor (VX-809) restores the ability of CF macrophages to phagocytose and kill Pseudomonas aeruginosa. American Journal of Physiology-Lung Cellular and Molecular Physiology 314: L432–L438, 2018. doi: 10.1152/ajplung.00461.2017.

58. Graeber SY, Renz DM, Stahl M, Pallenberg ST, Sommerburg O, Naehrlich L, Berges J, Dohna M, Ringshausen FC, Doellinger F, Vitzthum C, Röhmel J, Allomba C, Hämmerling S, Barth S, Rückes-Nilges C, Wielpütz MO, Hansen G, Vogel-Claussen J, Tümmler B, Mall MA, Dittrich A-M. Effects of Elexacaftor/Tezacaftor/Ivacaftor Therapy on Lung Clearance Index and Magnetic Resonance Imaging in Patients with Cystic Fibrosis and One or Two F508del Alleles. Am J Respir Crit Care Med Epub ahead of print, 2022. doi: 10.1164/rccm.202201-0219OC.

59. Carnovale V, Iacotucci P, Terlizzi V, Colangelo C, Ferrillo L, Pepe A, Francalanci M, Taccetti G, Buonaurio S, Celardo A, Salvadori L, Marsicovetere G, D’Andria M, Ferrara N, Salvatore D. Elexacaftor/Tezacaftor/Ivacaftor in Patients with Cystic Fibrosis Homozygous for the F508del Mutation and Advanced Lung Disease: A 48-Week Observational Study. J Clin Med 11: 1021, 2022. doi: 10.3390/jcm11041021.

60. Schnell A, Hober H, Kaiser N, Ruppel R, Geppert A, Tremel C, Sobel J, Plattner E, Woelfle J, Hoerning A. Elexacaftor - Tezacaftor - Ivacaftor treatment improves systemic infection parameters and Pseudomonas aeruginosa colonization rate in patients with cystic fibrosis a monocentric observational study. Heliyon 9: e15756, 2023. doi: 10.1016/j.heliyon.2023.e15756.

61. Sheikh S, Britt Jr. RD, Ryan-Wenger NA, Khan AQ, Lewis BW, Gushue C, Ozuna H, Jaganathan D, McCoy K, Kopp BT. Impact of elexacaftor–tezacaftor–ivacaftor on bacterial colonization and inflammatory responses in cystic fibrosis. Pediatric Pulmonology n/a, [date unknown]. doi: 10.1002/ppul.26261.

62. Miller AC, Harris LM, Cavanaugh JE, Abou Alaiwa M, Stoltz DA, Hornick DB, Polgreen PM. The Rapid Reduction of Infection-Related Visits and Antibiotic Use Among People With Cystic Fibrosis After Starting Elexacaftor-Tezacaftor-Ivacaftor. Clinical Infectious Diseases 75: 1115–1122, 2022. doi: 10.1093/cid/ciac117.

63. Cabak A, Hovold G, Petersson A-C, Ramstedt M, Påhlman LI. Activity of airway antimicrobial peptides against cystic fibrosis pathogens. Pathogens and Disease 78: ftaa048, 2020. doi: 10.1093/femspd/ftaa048.

64. Chen CI-U, Schaller-Bals S, Paul KP, Wahn U, Bals R. β-defensins and LL-37 in bronchoalveolar lavage fluid of patients with cystic fibrosis. Journal of Cystic Fibrosis 3: 45–50, 2004. doi: 10.1016/j.jcf.2003.12.008.

65. Tesse R, Cardinale F, Santostasi T, Polizzi A, Manca A, Mappa L, Iacoviello G, De Robertis F, Logrillo VP, Armenio L. Association of β-defensin-1 gene polymorphisms with Pseudomonas aeruginosa airway colonization in cystic fibrosis. Genes Immun 9: 57–60, 2008. doi: 10.1038/sj.gene.6364440.

66. Lavigne MC, Thakker P, Gunn J, Wong A, Miyashiro JS, Wasserman AM, Wei S-Q, Pelker JW, Kobayashi M, Eppihimer MJ. Human bronchial epithelial cells express and secrete MMP-12. Biochemical and Biophysical Research Communications 324: 534–546, 2004. doi: 10.1016/j.bbrc.2004.09.080.

67. Wagner CJ, Schultz C, Mall MA. Neutrophil elastase and matrix metalloproteinase 12 in cystic fibrosis lung disease. Mol Cell Pediatr 3: 25, 2016. doi: 10.1186/s40348-016-0053-7.

68. Churg A, Wang RD, Tai H, Wang X, Xie C, Dai J, Shapiro SD, Wright JL. Macrophage Metalloelastase Mediates Acute Cigarette Smoke–induced Inflammation via Tumor Necrosis Factor-α Release. Am J Respir Crit Care Med 167: 1083–1089, 2003. doi: 10.1164/rccm.200212-1396OC.

69. Jw P, Is S, Uh H, Sr O, Jh K, Ks A. Pathophysiological changes induced by Pseudomonas aeruginosa infection are involved in MMP-12 and MMP-13 upregulation in human carcinoma epithelial cells and a pneumonia mouse model. Infection and immunity 83, 2015. doi: 10.1128/IAI.00619-15.

70. Schönbeck U, Mach F, Libby P. Generation of biologically active IL-1 beta by matrix metalloproteinases: a novel caspase-1-independent pathway of IL-1 beta processing. J Immunol 161: 3340–3346, 1998.

71. Kassim SY, Gharib SA, Mecham BH, Birkland TP, Parks WC, McGuire JK. Individual matrix metalloproteinases control distinct transcriptional responses in airway epithelial cells infected with Pseudomonas aeruginosa. Infect Immun 75: 5640–5650, 2007. doi: 10.1128/IAI.00799-07.

72. López-Boado YS, Wilson CL, Parks WC. Regulation of matrilysin expression in airway epithelial cells by Pseudomonas aeruginosa flagellin. J Biol Chem 276: 41417–41423, 2001. doi: 10.1074/jbc.M107121200.

73. Manicone AM, Birkland TP, Lin M, Betsuyaku T, van Rooijen N, Lohi J, Keski-Oja J, Wang Y, Skerrett SJ, Parks WC. Epilysin (MMP-28) restrains early macrophage recruitment in Pseudomonas aeruginosa pneumonia. J Immunol 182: 3866–3876, 2009. doi: 10.4049/jimmunol.0713949.

74. McMahan RS, Birkland TP, Smigiel KS, Vandivort TC, Rohani MG, Manicone AM, McGuire JK, Gharib SA, Parks WC. Stromelysin-2 (MMP10) Moderates Inflammation by Controlling Macrophage Activation. J Immunol 197: 899–909, 2016. doi: 10.4049/jimmunol.1600502.

75. Lepissier A, Bonnel AS, Wizla N, Weiss L, Mittaine M, Bessaci K, Kerem E, Houdouin V, Reix P, Marguet C, Sermet-Gaudelus I, MODUL-CF Pediatric French Network for Cystic Fibrosis. Moving the Dial on Airway Inflammation in Response to Trikafta in Adolescents with Cystic Fibrosis. Am J Respir Crit Care Med 207: 792–795, 2023. doi: 10.1164/rccm.202210-1938LE.

